# Combinatorial CAR design improves target restriction

**DOI:** 10.1101/2020.10.09.333724

**Authors:** Hakan Köksal, Pierre Dillard, Asta Juzeniene, Gunnar Kvalheim, Erlend B. Smeland, June H. Myklebust, Else Marit Inderberg, Sébastien Wälchli

## Abstract

CAR T cells targeting the B-lymphocyte antigen CD19 have led to remarkable clinical results in B-cell leukemia and lymphoma, but eliminate all B-lineage cells, leading to increased susceptibility to severe infections. As malignant B cells will express either immunoglobulin (Ig) light chain κ or λ, we designed a second-generation CAR targeting Igκ, IGK CAR. This construct demonstrated high target specificity, but displayed reduced efficacy in the presence of serum IgG. Since CD19 CAR is insensitive to serum IgG, we designed various combinatorial CAR constructs in order to maintain the CD19 CAR T cell efficacy, but with IGK CAR target selectivity. The Kz-19BB design, combining CD19 CAR containing a 4-1BB co-stimulatory domain with an IGK CAR containing a CD3zeta stimulatory domain, maintained the target specificity of IgK CAR and was resistant to the presence of soluble IgG. Our results demonstrate that a combinatorial CAR approach can improve target selectivity and efficacy.

## Introduction

Chimeric antigen receptors (CARs) are engineered molecules which enable T cells to recognize and eliminate antigen positive target cells. The design of a CAR consists of the antigen recognition domain, usually derived from an antibody in which the variable fragments are arranged as a single-chain molecule (scFv) fused to different signaling domains (1). The first generation CARs contain a single signaling domain derived from the CD3 zeta (CD3ζ) protein, a crucial subunit of the CD3 complex which is involved in the early T-cell receptor (TCR) signaling, also known as signal 1. Due to the cellular exhaustion resulting from the use of this signal domain only, CARs were reinforced with a co-stimulatory signaling domain such as 4-1BB and/or CD28 in later generation constructs (a signal 2), which were then shown to improve proliferation and survival capacities of the CAR T cells (2, 3). It took 30 years for these molecules to be approved for clinical use, with the first therapeutic CAR target being CD19, a pan B-cell antigen, thus expressed in B-cell derived lymphomas and leukemias (4, 5). Numerous clinical studies demonstrated significantly improved outcomes in relapsed and refractory B-cell malignancies, and some of these studies were summarized in a recent review (6). As a ubiquitous marker of B cells, CD19 was an ideal antigen to limit on-target off-tumor toxicity but nonetheless resulted in complete B-cell aplasia (7-9). In a follicular lymphoma patient, Kochenderfer et al. demonstrated that complete B-cell eradication, subsequent to CD19 CAR treatment, resulted in a drastic decrease of serum Ig with increased susceptibility to infection. The low levels of serum Ig were compensated for by frequent intravenous Ig supplements to treat infectious diseases (10).

The B-cell receptor (BCR) is an attractive target as its expression is commonly maintained in malignant B cells (11). The BCR consists of two identical immunoglobulin (Ig) heavy and light chains, and due to allelic exclusion of immunoglobulin (Ig) genes, malignant B cells from an individual tumor are clonal for their BCR and express either Ig kappa (κ) or Ig lambda (λ) light chains (12, 13). It was previously observed that genetic deficiency of the κ-chain, resulting in a complete absence of κ^+^ IgG, did not prevent the patient from producing sufficient antibody titers to raise an immune response against infections (14). Thus, a κ^+^ B-cell aplasia can still be tolerable due to the presence of λ^+^ B cells. More than 10 years ago, an anti-Igκ (IGK) CAR construct was shown to be efficient in pre-clinical models (12) and later tested in a phase I clinical trial (15). However, a major issue was that T cells redirected with IGK CAR, although potent, were shown to be sensitive to Ig in serum (12), suggesting that the manufacturing of the therapeutic CAR T cells in the presence of human serum might affect the quality and the potency of the cells before injection.

In this study, we aimed at developing an IGK CAR resistant to soluble IgG (sIgG), in order to enhance B-cell malignancy treatment outcome by reducing CD19 CAR-related B-cell aplasia. We reasoned that splitting the stimulatory signal 1 and 2 of the CAR into an “AND” combinatorial system would result in CAR T cells with improved properties. This strategy is based on prior studies, which have shown that a combinatorial approach could be used to enforce tumor specificity (16). We identified the Kz-19BB CAR design, combining IGK-CD3zeta (Kz) with CD19-4-1BB CAR (19BB) CAR, as an improved CAR. It is Igκ-restricted and resistant to sIgG. We further show that CAR T cells achieved optimal activation only when both antigens, CD19 and Igκ, were present on the target cell, thus sparing the Igλ+ B cells. Hence, our combinatorial construct kept the advantages of both original CARs and additionally overcame their weaknesses. Our data support future development of “AND system” designs which can exploit different surface targets with an expression that is not entirely restricted to cancerous cells.

## Results

### IGK CAR T cells specifically kill Igκ+ target cells, but are inhibited in the presence of human serum and soluble IgG

We designed a second generation CAR by using the scFv from the anti-IGK hybridoma clone FN162 (Oslo University Hospital collection). To investigate the activity and specificity of the IGK CAR, we tested it against B-cell lymphoma cell lines with variable Igκ expression levels (REC-1, SU-DHL-4, BL-41, DAUDI and U2932), and included three Igλ+ cell lines (Mino, Granta-519, MAVER-1), one cell line which lacked Ig light chains (SC-1) and two non-B cell controls (Jurkat and K562) (**Fig. 1A and Fig. S1A**). We assessed the specificity of our IGK CAR construct by testing the cytotoxic activity of primary human T cells transfected with IGK CAR mRNA towards different cell lines and demonstrated high specificity (**Fig. 1B and Fig. S1B**). As the activity of IGK CAR might be inhibited in the presence of serum Ig, we next tested IGK CAR mediated killing of target cells in the presence of various dilutions of human serum. When comparing CD19 CAR and IGK CAR T cell efficacy against the Igκ+/CD19+ target cell line BL-41, we observed that killing efficacy of IGK CAR, unlike CD19 CAR, was markedly reduced in the presence of human serum (containing soluble IgG, IgA and IgM), even at dilutions as low as 3.12% (**Fig. 1C**). This was further confirmed when purified sIgG (50 μg/ml) was added to the IGK CAR T cell/target cell co-culture cytotoxicity assay, whereas the efficacy of CD19 CAR T cells remained unaltered (**Fig. 1D**).

**Figure 1:**
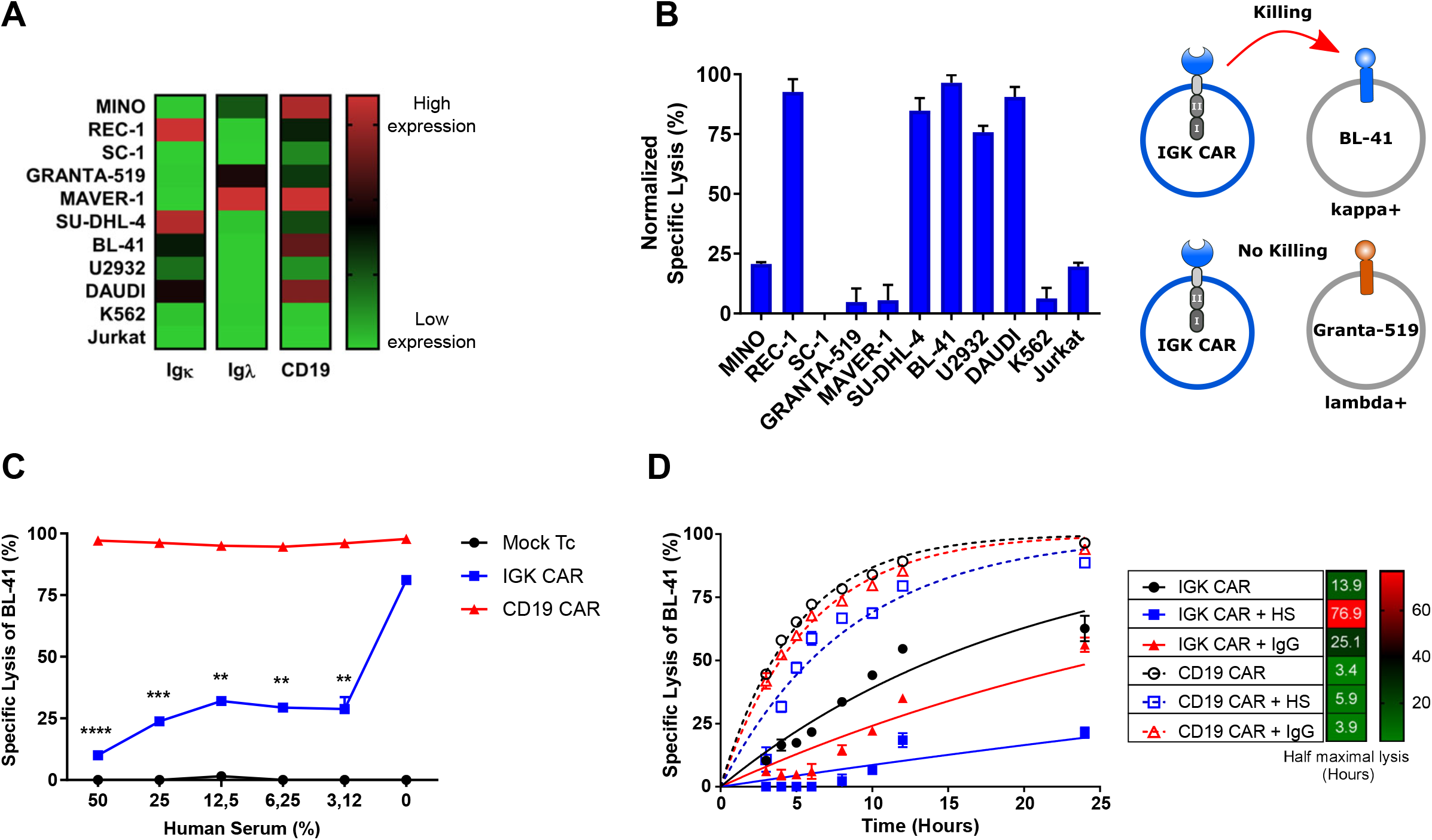
IGK CAR T cells target cancer cells specifically but are inhibited by human serum. (A) Flow cytometry analysis of Igκ, Ig λ light chains and CD19 expression on various cell lines. (B) BLI killing assay of Mock T cells and IGK CAR T cells against various target cells (25:1 E:T ratio). Shown is specific lysis as normalized values relative to Mock T cells after 6 hours of co-culture. Representative data from one of two independent experiments are shown. Data represent mean ± S.D. of quadruplicates. (C) BLI killing assay of Mock T cells, CD19 CAR and IGK CAR transfected T cells co-cultured with Igκ^+^ BL-41 lymphoma cell line for 10 hours (10:1 E:T ratio) in the presence of increasing human serum concentration. Data represent mean ± S.D. of triplicates. Representative data from one of two independent experiments are shown. (D) BLI killing assay of CD19 CAR and IGK CAR transfected T cells co-cultured with BL-41 cell line for 24 hours (10:1 E:T ratio) in the presence of 50% human serum or 50μg/ml of IgG. Half maximal lysis was obtained via one phase exponential fitting. Data represent mean ± S.D. of triplicates. Representative data from one of two experiments are shown. \*\**P* < 0.01, \*\**\*P* < 0.001, ****P<0.0001

### Combinatorial design of IGK-CD19 CAR overcomes soluble IgG sensitivity

To overcome the reduced efficacy of IGK CAR T cells in the presence of human serum whilst maintaining Igκ target restriction, we designed different “AND”-type constructs of CD19 or Igκ scFv using a single co-stimulatory domain, either 4-1BB domain (KBB and 19BB) or CD3ζ domain (Kz and 19z) (**Fig. 2A**). We first tested expression efficiency of these constructs in T cells after transient transfection alone or in combination (19z-KBB and Kz-19BB). We used two different detection methods (anti-Fab and protein-L) since recognition sensitivity varied among constructs. The combinatorial CAR constructs demonstrated similar expression levels as the single CARs (**Fig. 2B**).

**Figure 2:**
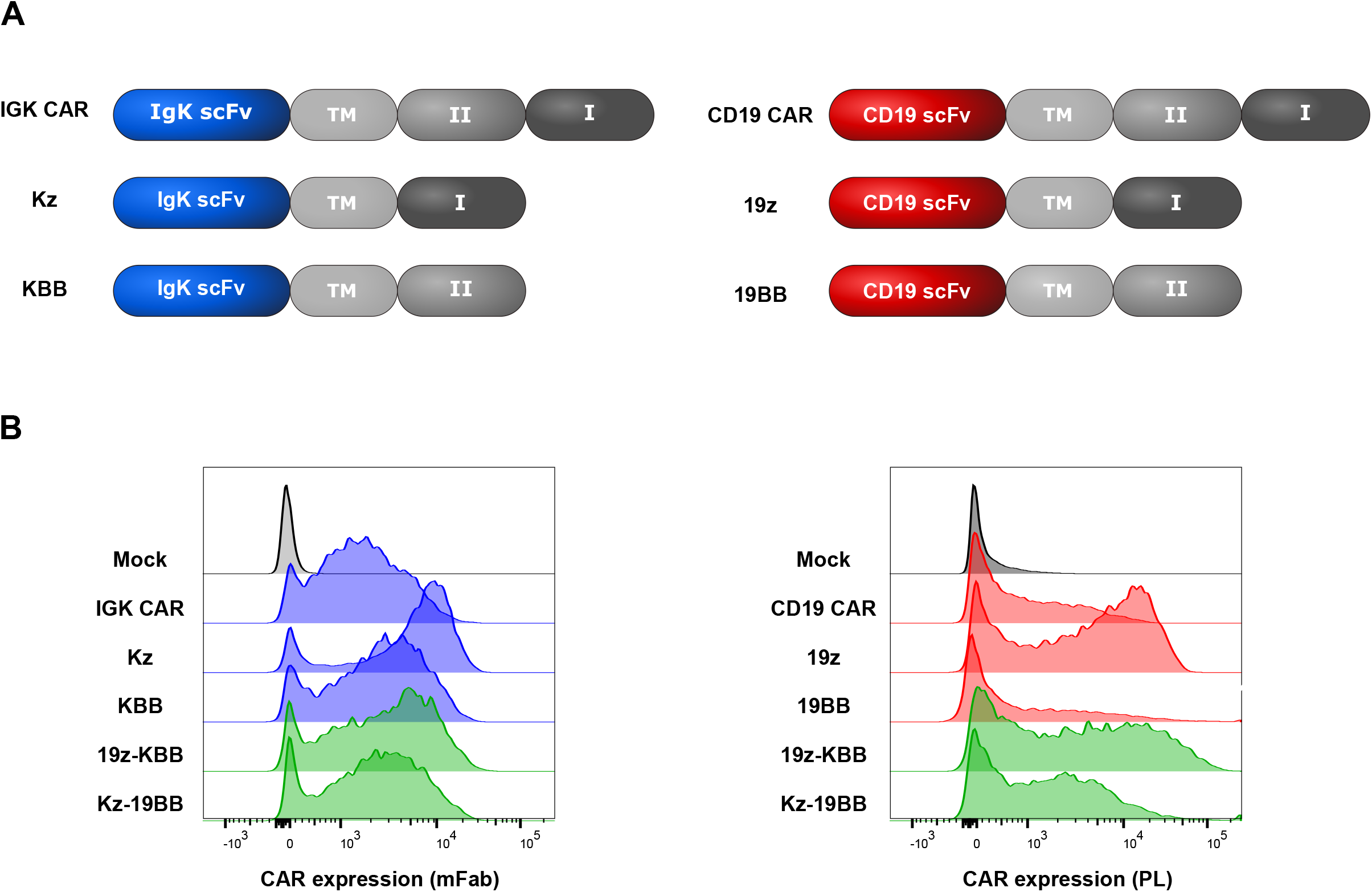
Design and expression of the classic and combinatorial CAR constructs. (A) Schematic representation of all classic and combinatorial CAR constructs.TM, CD8a transmembrane, I, CD3ζ, II, 4-1BB. (B) Igκ and CD19 scFv based mRNA constructs were electroporated into primary T cells. Igκ and CD19 redirected CAR expression was measured by staining with anti-mouse Fab and protein L, respectively. Representative FACS analysis of T cells 18 hours after electroporation is shown.

To test if the combinatorial CAR T cells could overcome inactivation by sIgG, we monitored the cytolytic activity of the different constructs against Igλ-/Igκ+/CD19+ BL-41 cells in presence or absence of sIgG (**Fig. S2A**). As shown in **Fig. 3A**, the IGK CAR T cells were sensitive to the presence of sIgG, whereas the Kz-19BB construct was insensitive. This suggests that the efficacy of CAR T cells was primarily regulated by the CD3ζ domain linked scFv and that the 4-1BB co-stimulatory domain by itself was not sufficient to induce significant cytotoxic activity. Furthermore, as expected, we did not observe any effect of sIgG when Granta-519 cells (Igλ+/Igκ-/CD19+) were targeted (**Fig. 3B**), supporting that Kz-19BB CAR had primarily acquired IGK CAR selectivity. All constructs alone or in combination were tested and confirmed that 19z-KBB CAR T construct followed CD19 CAR selectivity (**Figs. S2A and S2B**). These data further suggest that the Kz-19BB construct provided resistance to sIgG and confer the IGK CAR selectivity to Igκ+ cells. In order to quantify the sIgG inhibition, we also ran a killing assay with titrated soluble sIgG in the medium for all constructs, again demonstrating that Kz-19BB had acquired an advantage over IGK CAR (**Fig. 3C and Fig. S2C**).

**Figure 3:**
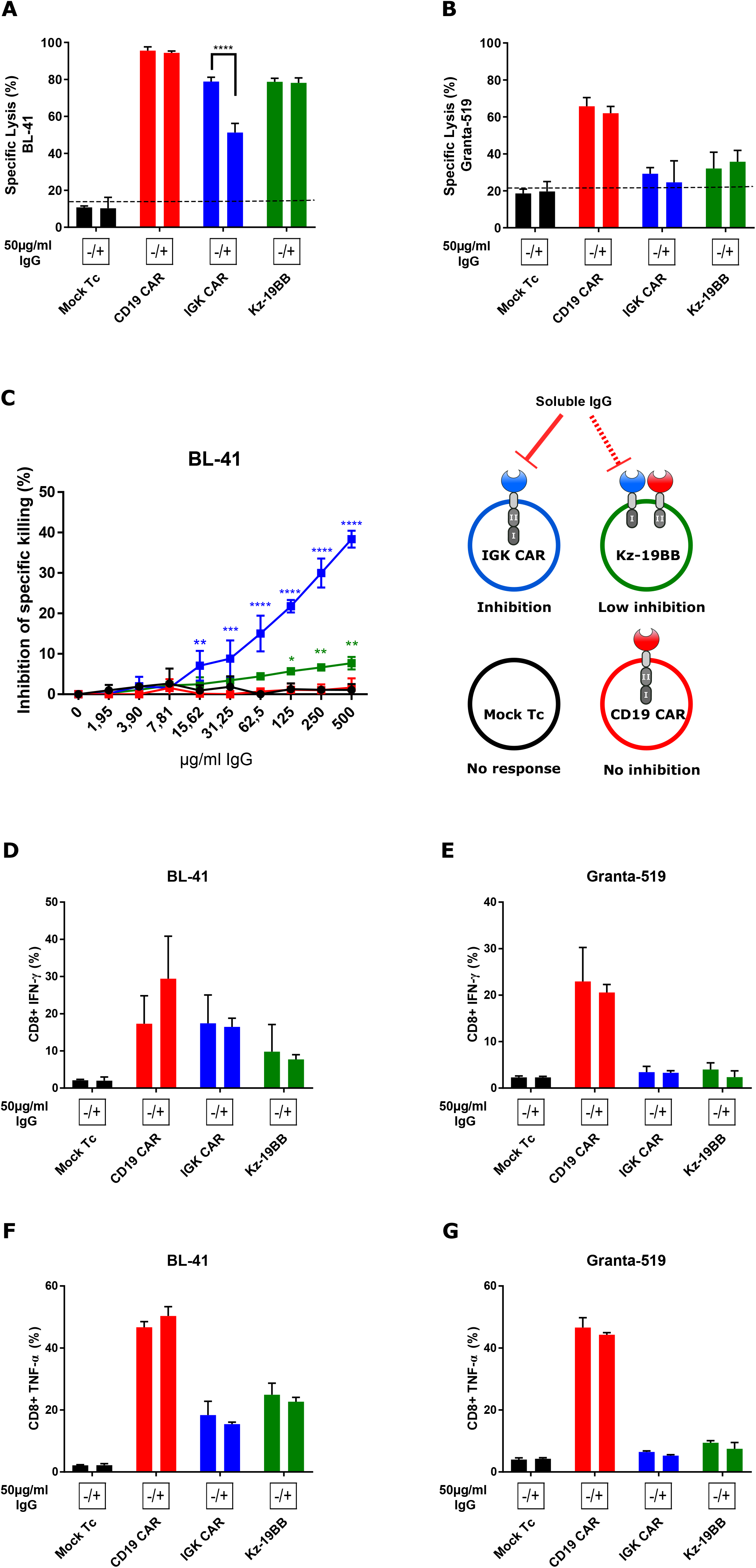
Combinatorial CAR T cells are less sensitive to serum inhibition while maintaining specificity. (A-B) BLI killing assay of Mock T cells and CAR construct electroporated T cells co-cultured with Igκ^+^ BL-41 or Igκ^-^ Granta-519 cell lines for 10 hours (10:1 E:T ratio) in the presence or absence of serum purified IgG (50μg/ml). Data represent mean ± S.D. of quadruplicates. Representative data from one of three independent experiments are shown. (C) BLI killing assay of Mock T cells and CAR construct electroporated T cells co-cultured with BL-41 cell line for 10 hours in presence of increasing concentration of IgG (10:1 E:T ratio). Specific lysis inhibition corresponds to the difference of cytotoxic capacity of each construct between IgG+ and IgG-conditions. Data represent mean ± S.D. of triplicates. One-way ANOVA was performed between Mock T cell and other groups for each IgG concentration. (D-G) CD8+ T-cell intracellular cytokine staining (IFN-γ and TNF-α) after co-cultivation with BL-41 or Granta-519 cell lines in the presence or absence of IgG at 50μg/ml (for 24 hours, 1:2 E:T ratio) Data represent mean ± S.D. of triplicates. Representative data from one of two independent experiments are shown. (A,B,D-G) Significance was assessed by Student t-test comparing IgG+ and IgG-conditions. \**P* < 0.05, \*\**P* < 0.01, \*\*\**P* < 0.001, ****P<0.0001.

We next analyzed cytokine production in the same conditions, where IGK CAR and Kz-19BB showed a reduced cytokine production compared to CD19 CAR expressing CD8 T cells in response to BL-41 cells, but were not sensitive to sIgG (**Figs. 3D-G**). The difference in sensitivity to sIgG in the two assays was probably due to the longer kinetics for cytokine response. These data were confirmed when all the constructs were run in combination or alone, with the exception of Kz alone being sensitive to sIgG in CD8 T cells (**Figs. S2D-G**) and CD4 T cells (**Figs. S3A-D**). This increased sensitivity could be due to the absence of a secondary co-stimulatory signal. As cytokine release was not affected by sIgG in our setting, these data confirm that selectivity of combinatorial constructs followed the identity of the CD3ζ domain carrying chain.

### Kz-19BB offers a trade-off between specificity and IgG insensitivity

We next studied the impact of the 19BB construct density on the cytotoxicity and on the IgG sensitivity of the Kz-19BB CAR (**Fig. 4 and Fig. S4**), using different concentrations of mRNA for electroporation to fine tune the surface density of the 19BB CAR constructs. First, we determined the baseline cytotoxic activity by Mock T cells with little to no activity (**Fig. 4A**). Then, we determined the baseline for IgG or HS sensitivity by IGK CAR and Kz which demonstrated substantial inhibition rates (**Figs. 4B and C**). However, the inhibition by serum or soluble Igs (sIg) was drastically decreased when Kz and 19BB were combined at a ratio of 1 to 0.5 (Kz-19BBx0.5) (**Fig. 4D**). At equimolar expression of Kz and 19BB or at increased concentrations of 19BB (Kz-19BBx2), the CAR T-cell mediated lysis of Igκ+/CD19+ BL41 target cells was even faster than with CD19 CAR and demonstrated potent killing even in the presence of IgG and HS (**Figs. 4E-H**). On the other hand, the same CAR T cells showed 19BB density-dependent cytotoxicity against Igλ+/CD19+ Granta-519 cells. Hence, higher 19BB concentration leads to a higher cytotoxic response towards the Granta-519 cell line (**Figs. S4A-F**). Even at the highest concentration of 19BB, the cytotoxic efficiency of combinatorial CARs against the Igκ-Granta-519 cell line was lower than CD19 CAR, demonstrating the conserved specificity to Igκ expressing targets of combined CARs (**Figs. S4F-H**). Furthermore, this suggests a correlation between the density of 19BB construct and the resistance to IgG inhibition, with specificity as a trade-off which is in line with previous studies, suggesting that the co-stimulatory signal improves the phosphorylation kinetics of the TCR signaling and boosts the response (17).

**Figure 4:**
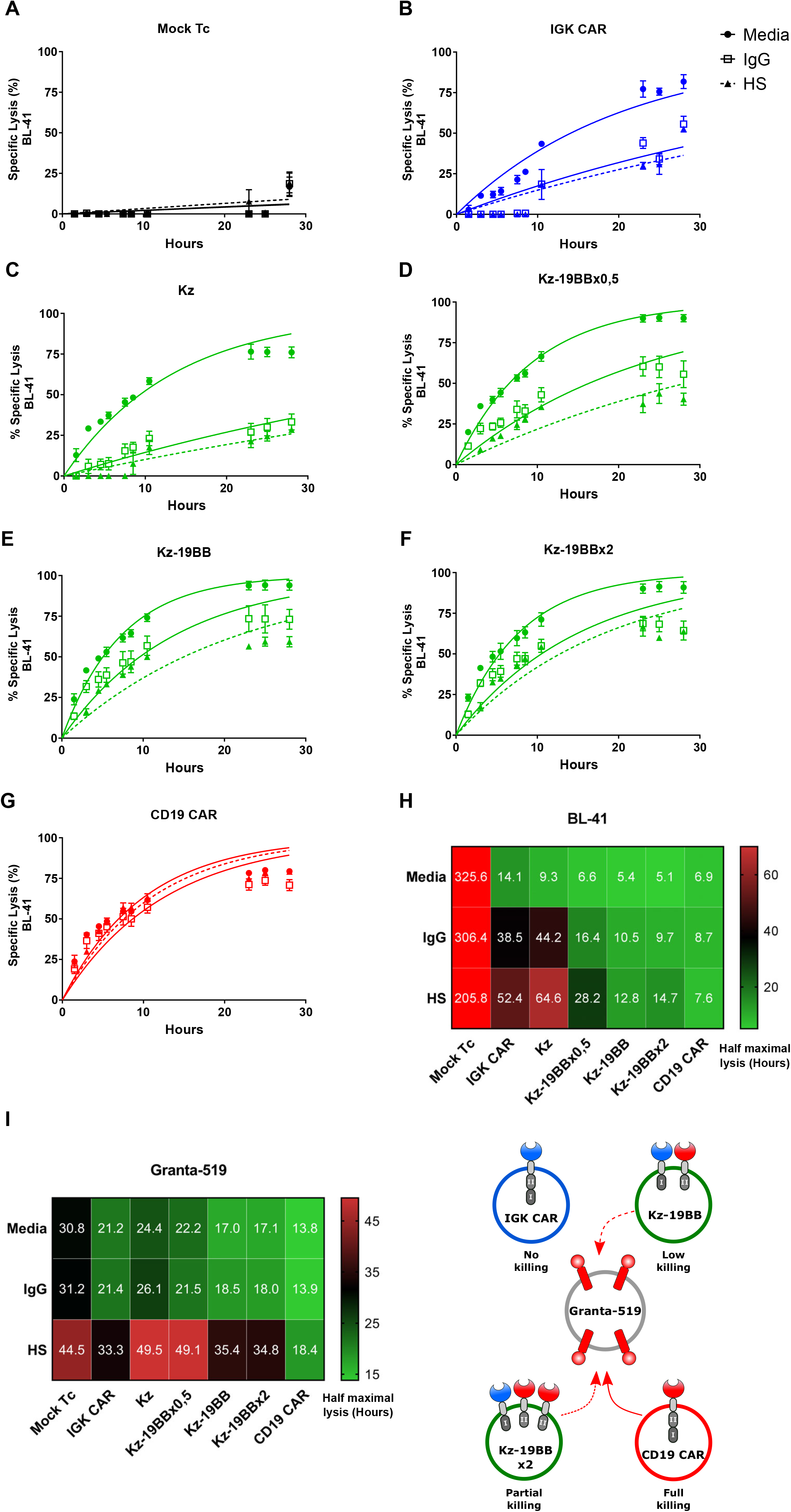
Kz-19BB offers a trade-off between IgG insensitivity and Igκ specificity. (A-G) BLI killing assay of Mock electroporated and CARs construct electroporated T cells co-cultured with BL-41 or Granta-519 target cell lines either in assay media, 50% human serum or 50μg/ml IgG (10:1 E:T ratio). Kz-19BB construct was prepared with varying concentrations of the 19BB part. Representative results only include analyses against BL-41 cell line. (H-I) Heat maps of half maximal lysis values obtained via one phase exponential fitting. Data represent mean ± S.D. of quadruplicates. Representative data from one of three experiments are shown. Significance was assessed by Student t-test comparing kappa+ and lambda+ cell numbers. \**P* < 0.05, \*\**P* < 0.01, \*\**\*P* < 0.001, ****P<0.0001.

### Kz-19BB conserves IGK CAR characteristics

To compare molecular and physiological characteristics of T cells expressing the combinatorial CARs, we monitored T cells electroporated with CAR constructs upon stimulation with surface coated antibodies where IgG was used as an Igκ specific stimulant and anti-CD3 as a general stimulant, independently of CAR specificity. Only IGK CAR and Kz-19BB T cells increased their metabolism (respiration capacity) upon incubation on an IgG-coated surface, exhibiting a state of immune activation. As expected, the same metabolic pattern and level of stimulation were observed when T cells were incubated on the anti-CD3 coated surface (**Figs. S5A-C**). In agreement with our previous data, 19z-KBB expressing T cells were not stimulated by IgG coating. We then studied the early signaling profile generated by the combined construct compared with the original ones. The activation profile of Kz-19BB CAR was measured by time-lapse imaging of ZAP-70 kinase phosphorylation (**Fig. S5D**). A synthetic surface was obtained after glass-coating of IgG, anti-CD3 or PLL. Similar to IGK CAR T cells, the combinatorial Kz-19BB CAR T cells were responsive to IgG and anti-CD3 coating by transient phosphorylation of ZAP-70 kinase. In contrast, CD19 CAR showed only anti-CD3 induced phosphorylation (**Fig. S5E**). These results were confirmed by measuring the density of adhered cells per unit of area, a direct measurement of T-cell activation (**Fig. S5F**). Taken together, these results demonstrated a comparable activation phenotype of Kz-19BB and IGK CAR cells upon Ig-kappa target.

### Impact of sIgG on Igκ targeting CAR T cells

We next studied if sIgG had an effect on CAR T-cell expansion. In order to use a system close to common clinical manufacturing, we employed a retroviral expression system and designed a combinatorial retroviral construct where Kz and 19BB were separated by a 2A ribosome skipping sequence which guarantees a close to equimolar production of the two CARs (18). Three separated constructs encoding IGK CAR, Kz-19BB CAR and CD19 CAR were prepared (**Fig. 5A)** and expressed in T cells (**Fig. 5B**). These cells were expanded into two separate cultures either with human serum (HS) or with serum replacement (SR) for 11 days and the cell number, the viability and the level of CAR expressions were monitored. Overall, T cells expanded more efficiently in HS than in SR. We also observed that IGK CAR T cells were slightly less confluent than all other HS expanded T cells (**Figs. 5C and 5D**), which could be reminiscent of this constant stimulation of the receptor by sIgG. We next analyzed the cell viability and, in agreement with previous observations, we noticed that HS expanded cells overall led to more viable cells than SR expanded cells and detected no difference between IGK CAR and Kz-19BB expressing T cells (**Fig. S6A**). To investigate the general trend, we divided SR values over HS (**Fig. S6B).** Finally, the expression of IGK CAR and Kz-19BB T cells was decreased when expanded in HS whereas CD19 CAR T cells were not affected (**Figs. S6C-F**), thus Kz-19BB could not counteract the sIgG effect after long term stimulation. Again this effect was probably linked to a constitutive recycling of a stimulated receptor. Importantly, although the percentage of CAR expression was not dramatically affected (**Figs. S6C and S6D**) the intensity of CAR expression was decreased by 3-fold at the end of the expansion (**Figs. S6E and S6F**). In line with these observations, we were able to detect that IGK CAR and Kz-19BB were stimulated and expanded by incubation with soluble or coated IgG (**Figs. S7A and B**). These last data are in agreement with studies showing the stimulating effect of dimeric soluble factors on CAR T cells (19). Together, these data point toward a mild improvement of Kz-19BB over IGK CAR in HS containing culture medium, in terms of cell number. However, the loss of CAR expression and the viability could not be completely overcome.

**Figure 5:**
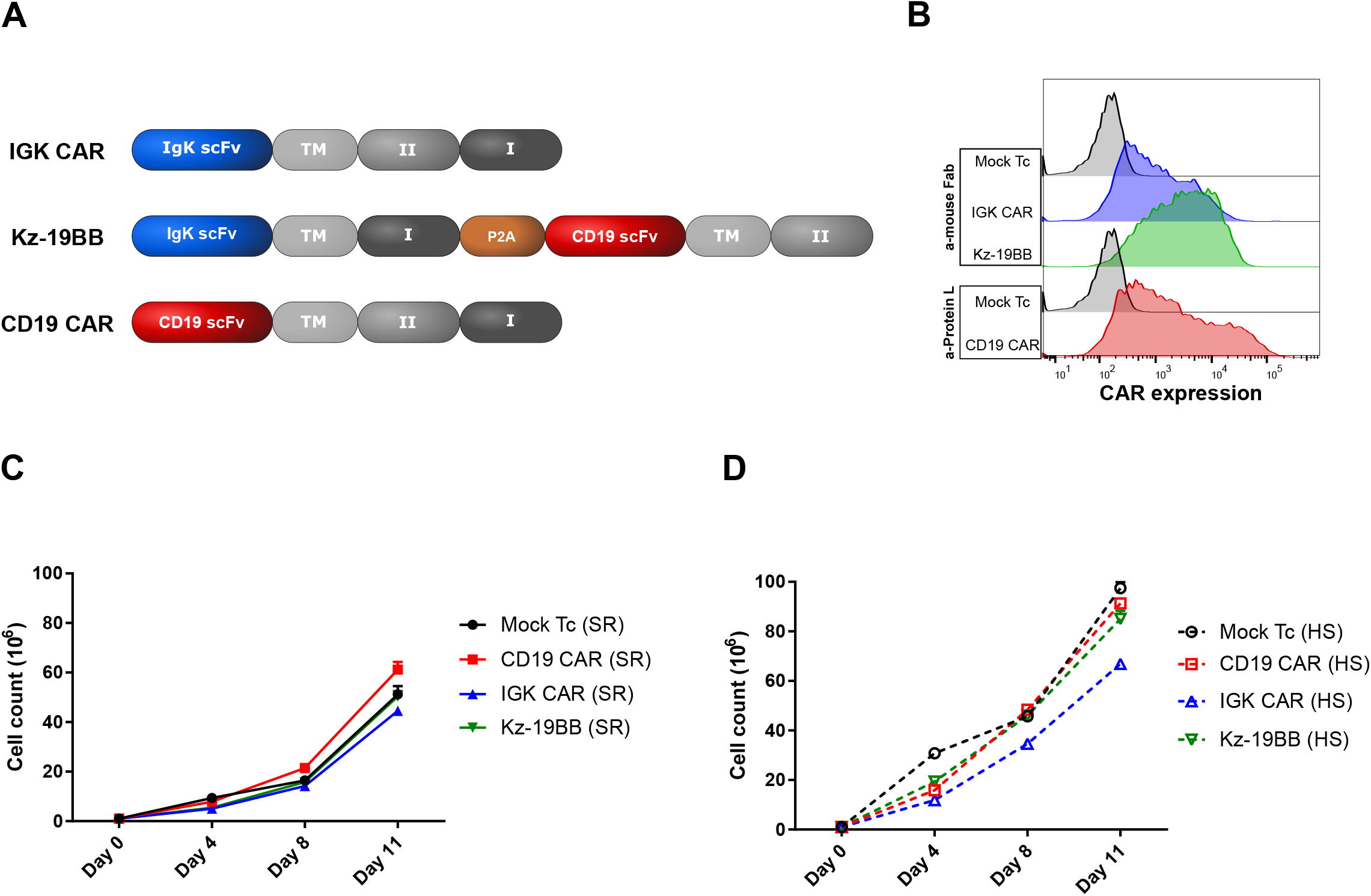
Impact of human serum on stable IGK CAR-expressing T cell expansion. (A) Schematic representation of Kz-19BB, IGK and CD19 CAR constructs. (B) CAR expressions were analyzed after transduction and 11 days expansion of primary T cells by staining with anti-mouse Fab or Protein L antibody. (C-D) Mock and CAR expressing T cells were expanded in either SR (serum replacement) or HS (human serum) containing X-VIVO-15 complete medium for 11 days. Cell counts were evaluated on days 0, 4, 8 and 11 by a Countess cell counter utilizing trypan blue exclusion.

### Stable Kz-19BB expressing T cells demonstrate IGK CAR selectivity

We then tested whether stable equimolar expression maintained Igκ selectivity. Specificity and cytotoxic capacities of the transduced T cells were assessed in a BLI assay against BL-41 and Granta-519 cell lines, with similar results to what was observed upon mRNA electroporation (**Figs. 6A and B**). However this statement might not apply to a situation of high CAR expression or saturated antigen density (see discussion). We further verified the selectivity of Kz-19BB CAR in a mixture of target cells (Igκ+ and Igκ-). As shown, unlike CD19 CAR T cells, IGK CAR and Kz-19BB CAR T cells selectively killed Igκ+/CD19+ BL-41 cells but not Igκ-/CD19+ Granta-519 (**Figs. 6C-E**). We confirmed these observations using Epstein Barr virus (EBV) transformed primary B cells co-cultured with IGK and Kz-19BB CAR T cells and demonstrated that Igκ+ EBV+ B cells were specifically eliminated whereas CD19 CAR eliminated all B cells (**Figs. S8A-C**). We finally confirmed the target restriction of our construct by including an osteosarcoma cell line OHS (CD19-/Igκ-) (**Figs. S8D and E**). These data suggest that the scFv linked to the CD3ζ domain determined the target selectivity whereas the scFv linked to the 4-1BB domain could potentiate the efficacy of the T cells upon expression of the combinatorial CAR.

**Figure 6:**
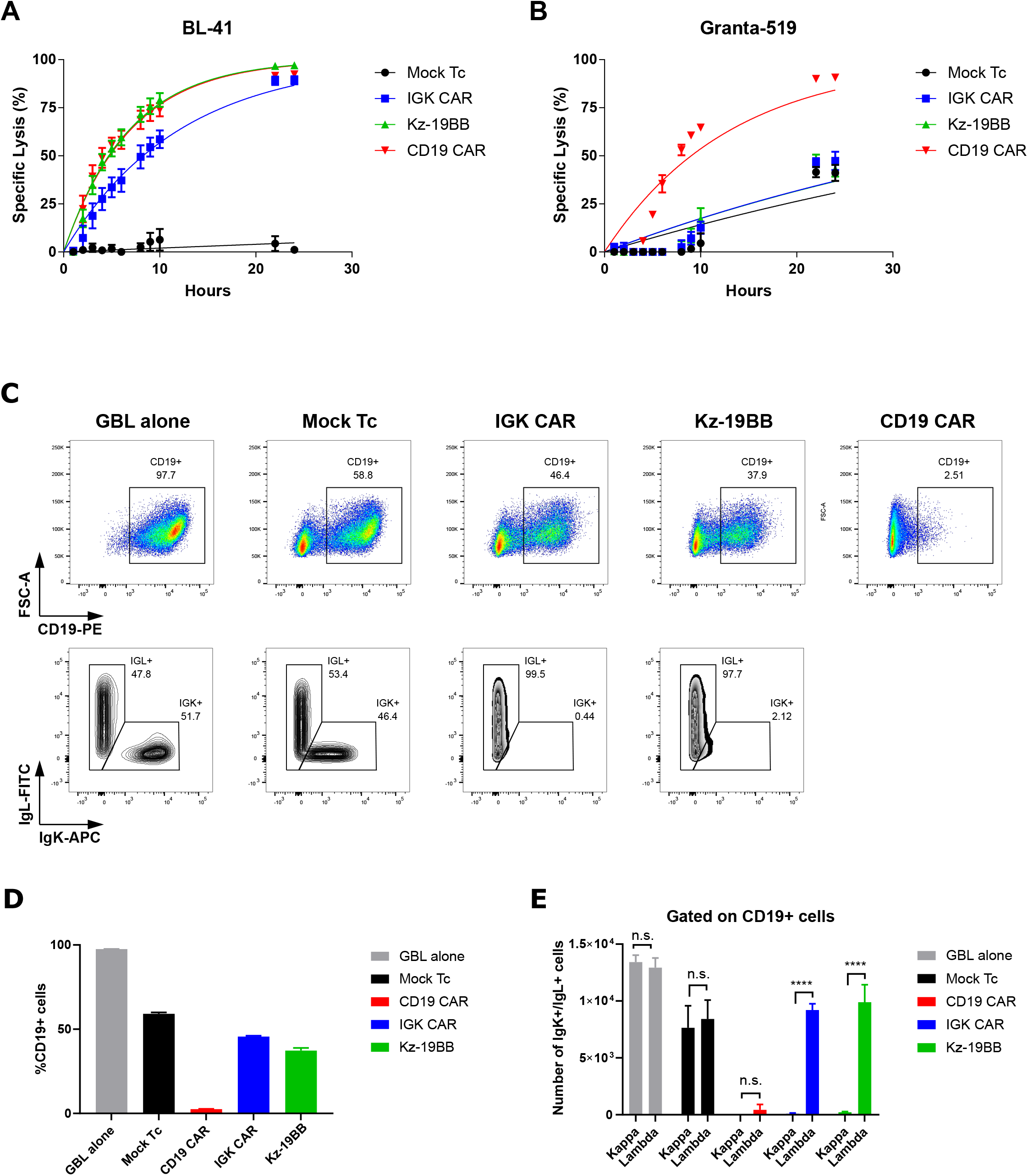
Kz-19BB is efficient when stably expressed and demonstrates similar specificity against mixed cell cultures. (A-B) BLI killing assay of Mock and CAR transduced T cells co-cultured with BL-41 or Granta-519 target cell lines (10:1 E:T ratio). Data represent mean ± S.D. of triplicates. Representative data from one of two experiments is shown. (C-E) Retrovirally transduced T cells co-cultured for 12 hours with both BL-41 and Granta-519 target cell lines at a ratio of 2:1:1, respectively. After co-culture cells were stained with anti-CD19-PE, anti-Igκ APC and anti-Igλ-FITC. Data represent mean ± S.D. of quadruplicates. Data pooled from two independent experiments. \**P* < 0.05, \*\**P* < 0.01, \*\*\**P* < 0.001, ****P<0.0001.

### Kz-19BB is efficient against 3D tumor spheroids

We finally assessed the efficacy of Kz-19BB against 3D spheroid tumors and compared it to the performance of the single CARs. We established 3D lymphoma spheroids to analyze the efficacy and specificity of Kz-19BB against tumor formations that are related to *in vivo* structures. To this end, BL-41 and Granta-519 spheroids were prepared on agar coated wells. T cells were added when spheroid diameters were around 1 μm. Annexin V substrate was used to monitor apoptosis by live cell imaging. In agreement with the killing assays, BL-41 spheroids were lysed by all CAR T cells (**Figs. 7A and B**) whereas only CD19 CAR T cells demonstrated significant impairment of Granta-519 spheroid growth (**Figs 7C and D**). Together these data confirm that combinatorial the Kz-19BB CAR construct was as efficient and selective as the original second generation IGK CAR in controlling tumor growth in a complex tumor structure.

**Figure 7:**
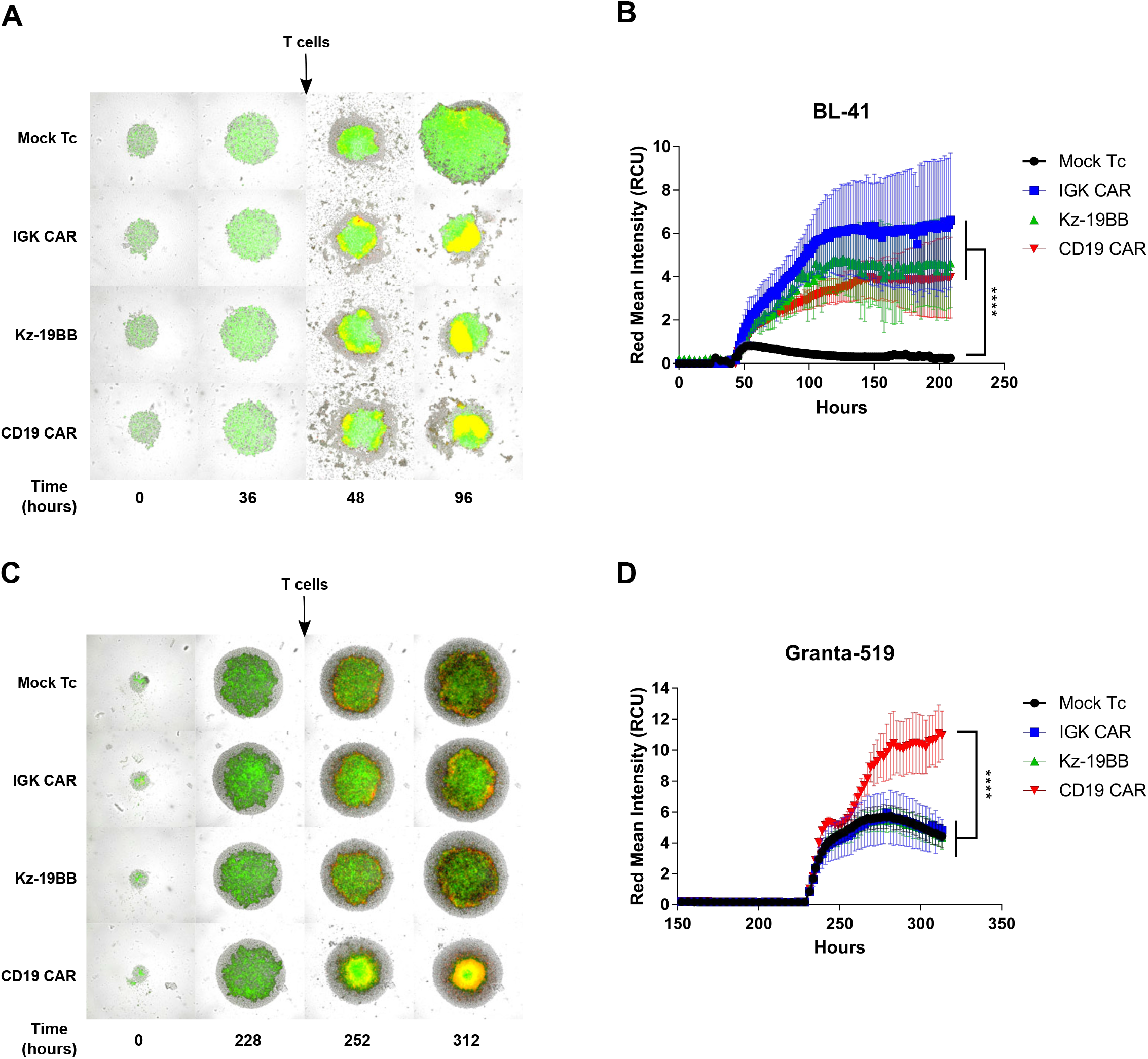
Kz-19BB is efficient when stably expressed and demonstrates similar specific potency against 3D tumor cultures. (A, C) Representative micrographs of BL-41 and Granta-519 spheroids, respectively, co-cultured with either Mock, IGK CAR, Kz-19BB or CD19 CAR expressing T cells. Both cell lines are GFP/Luc+. Red signal represents the presence of Annexin V. (B, D) Annexin V measurements of spheroids over time. Data represent means ± SD of hexaplicates. Representative data from one of two independent experiments are shown. Two-way ANOVA test was performed to compare groups. ****P<0.0001.

## Discussion

CD19 CAR T-cell therapy has shown remarkable clinical efficacy in multiple subtypes of B-cell lymphoma (20) and led to the FDA approval of two CD19 CAR products for relapsed or refractory B-ALL and aggressive non-Hodgkin lymphoma treatment (21-24). However, this treatment comes with a critical pitfall: by directing T cells towards the CD19 antigen, a common lineage marker, the entire B-cell population is indiscriminately eradicated which in turn impairs patients’ humoral immune response (11). There is therefore a need for alternative target antigens with less off-tumor toxicity. To this end, we and others (12) selected a restricted lineage marker, the Igκ light chain of the BCR. The clonal light chain expression on most B-NHL subtypes makes this an ideal therapeutic target to limit on-target, off-tumor toxicity. Despite some objective responses demonstrated after treatment with IGK CAR in a phase I clinical trial (15) where the impact of soluble Igκ was discussed, no data showing the resistance to sIgG were presented. We reasoned that despite its serum sensitivity, IGK CAR could improve CD19 CAR construct selectivity in a combinatorial “AND” format (16). Since this format requires splitting of the signaling unit, T-cell sensitivity to serum would be attenuated due to the lack of co-stimulation provided by the CD19 CAR in the presence of serum proteins.

The initial evaluation of our IGK CAR demonstrated that it was potent and specific, but sensitive to human serum and sIg, thus in agreement with previous reports (12). We combined IGK CAR and CD19 CAR with different signaling domains and confirmed previous reports showing that the CD3ζ domain (z) was the major driver of the CAR T cell activation determining the specificity of the construct. As expected, the co-stimulatory domain 4-1BB (BB) was not able to trigger a significant CAR-mediated T cell killing by itself but could be exploited to attenuate serum sensitivity. Indeed, even if the adjunction of the 19BB construct to the Kz part did not boost the activation, it drastically reduced the inhibition caused by human serum. Further analyses of the Kz-19BB combination demonstrated similarity with IGK CAR in terms of specificity and activation (in response to IgG or Igκ ligand binding). For these experiments, we used mRNA co-electroporation to adjust individual components and hence control the protein levels. Besides flexibility regarding specificity and sensitivity towards IgG inhibition, the mRNA concentration had a major impact on the functional outcome. This was in line with a recent publication demonstrating that a two-output biological circuit can be pushed in one direction or the other by altering mRNA concentrations alone (25). Accordingly, we used varying concentrations of the 19BB part and observed that design could be even more intricate and flexible than intended. Each increasing concentration of 19BB yielded a CAR construct less sensitive to IgG inhibition, but with specificity closer to that of CD19 CAR, allowing us to control the trade-off. This concept provides additional advantages such as the interchangeability of the 4-1BB linked parts. As an example, in the case of CD19 negative relapse, one could use an alternative targeting scFv, such as CD22 (26) or CD37 (27, 28) to overcome the loss of CD19.

We also assessed the efficacy and specificity in retroviral settings. Including a P2A ribosomal skipping sequence allowed us to have an equimolar expression of Kz and 19BB parts. During the expansion comparison of the T cells, we observed that HS provides a better T-cell environment leading tohigher expansion rates and higher viabilities than SR. Although a slight benefit of Kz-19BB was detected in the cell number after 11 days of expansion, both IGK and Kz-19BB CAR T cells demonstrated lower CAR expression in HS. This suggests that both activation antibodies and serum components in the media are over-stimulating IGK and Kz-19BB constructs leading to elimination of high intensity CAR expressing T cells or important recycling. Thus the combinatorial CAR, although efficient in maintaining selective killing, was not able to overcome the negative effect of the sIgG on the CAR T cells. It is tempting to speculate that this was due to normal receptor recycling upon ligand binding, but whether this will impact the final clinical product will need to be evaluated. Nevertheless, this issue is manageable; one could either use SR instead of HS or alternative T-cell media which do not require serum supplementation (29).

In order to proceed with the evaluation of Kz-19BB, we performed 3D tumor spheroid killing assays because they mimic some of the *in vivo* tumor properties (30, 31). Despite the expansion downsides, both IGK CAR and Kz-19BB still demonstrated significant cytotoxicity against BL-41 spheroids and maintained their selectivity against Granta-519 spheroids.

The construct presented herein is a prototype and alternative versions will be designed in which the affinity of one of the scFv can be modified: it is tempting to speculate that a lower affinity CD19 CAR (32) would reduce the recognition of Igλ+/CD19+ targets. A similar observation was also noted with our high affinity anti-Igλ (IGL) CAR (33) when designed in a CD19-combinatorial construct (unpublished data). Another possible improvement would be the regulation of the expression: a lower or controlled presence of the construct could also affect the Igλ+/CD19+ target recognition as supported by our mRNA titration studies, where increase IGK part reduced CD19 CAR dominancy. To overcome the variable CAR expression, one could CRISPR guide the combinatorial CAR to have a predictable and comparable expression level in each transduction (34).

In summary, we describe an alternative CD19 CAR which becomes selective through IGK combination to avoid B-cell aplasia. One can predict that this format will be used to combine alternative targets, thus improving selectivity, which should result in an increased safety.

## Experimental Procedures

### Plasmid design

A DNA sequence encoding the anti-Igκ scFv was generated after sequencing the original hybridoma (FN162). Briefly, the sequences of the VL and VH regions were determined by 5’-RACE (33). The design consists in the linkage of the two chains with a (G_4_S)_4_ linker. Synthetic sequences were acquired at Eurofins MWG (Ebesberg Germany). The scFv for the CD19 CAR (fmc63 clone) was a kind gift from Martin Pule (University College London, UK) that we subcloned in our codon optimized second generation signaling tail which is composed of CD8 hinge and transmembrane domain linked to 4-1BB and CD3ζ. The scFvs and the signaling tail were subcloned into pENTR Gateway (Themofisher, Waltham, MA, USA) and further subcloned in compatible expression vectors (35). Combinatorial CAR clonings were performed by modified CAR tails. The constructs only containing 4-1BB were prepared by introducing a stop codon via site-directed mutagenesis after 4-1BB domain with the following primers (5’-3’); forward GGTTGTGAGCTGTGAGTGAAGTTTTCC, reverse GGAAAACTTCACTCACAGCTCA CAACC. The constructs with only CD3ζ tail were synthesized by Eurofins-MWG and fused to the scFv sequences. To clone Kz-2A-19BB construct for retroviral expression, the sequence coding for a partial site from the end of Igκ scFv and rest of the CAR tail, P2A ribosome skipping sequence ^26^ and partially the beginning of CD19 scFv sequence were synthesized by Eurofins-MWG. The ligation of different element of the final construct was performed into a pENTR vector. The firefly luciferase-GFP fusion protein coding sequence (a kind gift from Rainer Löw, EUFETS AG, Germany) (36) was incorporated into pMP71 and used to stably transduce target cell lines as reported in (37).

### Cell lines, media

The human cell lines, BL-41, GRANTA-519, DAUDI, REC-1, SU-DHL-4, U2932, SC-1, MINO, K562 and MAVER-1 were obtained from DSMZ. J76 was a kind gift from M. Heemskerk (Leiden University Medical Center, The Netherlands). EBV transformed B cell lines were generated in house. The cells were cultured in RPMI 1640 (PAA, Paschung, Austria) and supplemented with 10% fetal calf serum (FCS, PAA) and 50 μg/mL Gentamycin (Thermo Fischer, Oslo, Norway). Phoenix-AMPHO (CRL-3213) cell line was purchased from ATCC, Manassas, USA and was maintained in DMEM (Sigma-Aldrich, Oslo, Norway) supplemented with 10% FCS and 50 μg/mL Gentamycin.

### Soluble IgG purification

Soluble IgG (sIgG) were purified directly from human serum using Pierce™ Protein A/G Agarose (Thermo Fischer Scientific) by following the manufacturer’s protocol. In brief, first Protein A/G agarose was loaded on Micro Bio-Spin™ Chromatography Columns (Bio-Rad Laboratories, Hercules, CA, USA). After gravity flow, human serum was loaded. sIgG was eluted from the agarose by 0.1 M Glycine pH 2, then neutralized by 1 M Tris pH 8. The concentration of purified IgG was measured on Nanodrop (Thermo Fischer Scientific).

### Synthetic mRNA preparation

*In vitro* transcribed (IVT) mRNA was synthesized using RiboMAX T7 Kit (Promega, Madison, WI, USA) as described (37, 38). Anti-Reverse Cap Analog (ARCA, Trilink Biotechnologies, San Diego, USA) was used for mRNA capping. The IVT mRNAs were evaluated by agarose gel electrophoresis and Nanodrop (Thermo Fischer Scientific, Waltham, USA) for quality and quantity, respectively.

### *In vitro* expansion of human T cells

Human PBMCs were isolated from healthy donors by a protocol adapted from T-cell production under GMP conditions as previously described (39). In brief, PBMCs were separated from blood by density gradient and cultured in the presence of Dynabeads (Dynabeads^®^ *ClinExVivo™* CD3/CD28, ThermoFischer, Oslo, Norway) in X-VIVO 15 (Lonza, Basel, Switzerland) supplemented with 100U/mL recombinant human IL-2 (Proleukin, Prometheus Laboratories Inc., San Diego, USA) and 5% human serum for 11 days. When specifically mentioned CTS serum replacement (Thermo Fischer Scientific) was used instead of human serum. On specific days, a small volume of media was separated from culture and subjected to Countess™ II Automated Cell Counter (Thermo Fischer Scientific) to monitor viability and numbers during the expansion phase. Expanded T cells were frozen in batches and stored in liquid nitrogen for future manipulation.

### IVT mRNA electroporation of human T cells

Expanded T cells were washed twice with non-supplemented RPMI media and resuspended at 70×10^6^ cells/mL. The IVT mRNA was mixed with the washed T cells at a concentration of 100 μg/mL. The mix was transferred to a 4-mm cuvette and electroporated at 500 V and 2 ms using a BTX 830 Square Wave Electroporator (BTX Technologies Inc., Hawthorne, USA). After electroporation T cells were transferred to complete culture medium and then left at 37 °C in 5% CO_2_ overnight. In co-electroporation cases, the total concentration of IVT mRNA for each construct was halved. In IVT mRNA titration case, the total mRNA concentration was maintained by an irrelevant GFP coding mRNA.

### Retroviral transduction of human T cells

Retroviral supernatants were collected as previously described (25). PBMCs were isolated from healthy donors as described above. After isolation, PBMCs were resuspended in complete X-VIVO 15 medium and transferred to a 24-well plate coated with anti-CD3 (1 μg/mL, OKT clone, ThermoFischer, Norway) and -CD28 (1 μg/mL, CD28.6 clone, ThermoFischer, Norway) at 1×10^6^ cells/well. Cells were then left at 37 °C in 5% CO_2_ for 3 days. Activated T cells were transferred to another 24-well plate (Nunc A/S, Roskilde, Denmark), pre-coated with retronectin (50 μg/mL, Takara Bio. Inc., Shiga, Japan). Activated T cells were spinoculated at 32 °C at 750 g for 60 minutes. The spinoculation was repeated once more the following day with fresh medium and retroviral supernatant. Afterwards, T cells were washed twice with complete X-VIVO 15 culture medium, transferred to a new 24-well plate and maintained in culture for transduction efficiency assessment. Transduced T cells were expanded with anti-CD3/28 Dynabeads as described above. Expanded T cells were frozen in aliquots and transferred to liquid nitrogen for future use.

### Functional assay and flow cytometry

Expression validations were done by flow cytometry. Cell lines or primary cells were washed twice with flow buffer (PBS with 2% FCS). Cells were then resuspended in antibody containing flow buffer for 15 minutes at room temperature and washed twice with flow buffer. Validation of CAR expression was performed with two antibodies. All CAR combinations with anti-Igκ scFv were assessed with anti-mouse Fab (Biotin-SP (long spacer) AffiniPure F(ab’)_2_ Fragment, Jackson ImmunoResearch, Cambridgeshire, UK) and CAR combinations with anti-CD19(fmc63) scFv were assessed by staining with Protein-L (Biotin-Protein L, GenScript, Piscataway, NJ, USA). In both cases, a Streptavidin-PE antibody (BD Biosciences, Franklin Lakes, NJ, USA) was used as a secondary antibody. Additionally, the following antibodies were used in the marker evaluation of cell lines; CD19-PE (BD Biosciences), Ig light chain κ-APC, and Ig light chain λ-PE (Biolegend, San Diego, USA).

T cells were electroporated as described above and cells were maintained in culture for 18 hours. CAR-expressing T cells were then co-cultured with the target cells at an E:T ratio of 1:2 for 6 hours. Culture media consists of X-VIVO 15 medium containing brefeldin A (Golgi-Plug, BD Biosciences) and monensin (Golgi Stop, BD Biosciences). Cells were then stained both for extracellular and intracellular markers using the PerFix-nc kit following the manufacturer’s protocol (Beckman Coulter, Indianapolis, USA). The following antibodies were used during the staining protocol; CD4-BV421 (Biolegend), CD8-PeCy7, IFNγ-FITC (eBiosciences, ThermoFischer), TNFα-PE (BD Biosciences, USA). Cells were acquired using a BD FACSCanto flow cytometer and the data were analyzed by Flow Jo software (Treestar Inc., Ashland, USA).

### Bioluminescence (BLI) cytotoxic assay

The protocol was previously described in detail (37). In brief, luciferase expressing target cells were transferred to a white round-bottomed 96-well plate in the presence of Xenolight D-Luciferin (75 μg/mL; Perkin Elmer, Oslo, Norway). CAR-expressing T cells were subsequently added in the mix and cells were cultured in an incubator (37 °C in 5% CO_2_) for the duration of the assay. When indicated, additional supplements such as sIgG and serum were added in the CAR-expressing T cells mix. Luminescence was monitored at every time point by a luminometer (Victor Multilabel Plate Reader, Perkin Elmer) as relative light units (RLU). For each assay one group of target cells were cultured alone to determine baseline lysis and another group was cultured in 1% Triton™ X-100 (Sigma-Aldrich) to determine the maximum lysis. Percentage lysis was calculated with the following equation; % specific lysis = 100x(spontaneous cell death RLU-sample RLU)/(spontaneous death RLU – maximal killing RLU).

### ZAP-70 phosphorylation

Glass slides (Ibidi, Germany) were washed with boiling piranha solution (70% of pure sulphuric acid with 30% of a 30% hydrogen peroxide solution) for 30 min. After intensive washing with PBS, glass slides were functionalized with 5 μg/mL of Anti-CD3 (OKT3) (eBiosciences, Thermo Fisher Scientific) or 5 μg/mL of IgG or 100 μg/mL of PLL for 30 minutes at room temperature followed by extensive rinsing with PBS + 0.1% of BSA. 8×10^5^ electroporated T-cells were washed and resuspended in PBS + 0.1% of BSA, then incubated on each substrate and fixed with 4% of Paraformaldehyde during 20 min (Sigma, Germany) after 2, 4, 8, or 16 min after incubation. Cells were washed with PBS + 0.1% of BSA and incubated in 50mM of NH_4_Cl (Sigma, Germany) for 20 min and then rinsed again PBS + 0.1% of BSA. Cells were permeabilized with 0,5% of Triton-X 100 in PBS + 0.1% of BSA during 15 min before being rinsed with PBS + 0.1% of BSA. Cells were then labelled with 0.1 μg/mL of Phospho-ZAP70/Syk (Tyr319, Tyr352) Monoclonal Antibody (n3kobu5) PE (eBioscience, Thermo Fisher Scientific) in PBS + 0.1% of BSA + 0.05% of saponin (Sigma, Germany) during 1 hour. After extensive rinsing with PBS, cells were imaged using a confocal microscope (Zeiss LSM 880 AiryScan) with a 63x 1.4 NA objective.

### SeaHorse mitostress assay

A Seahorse Extracellular Flux (XF96e) Analyzer (Agilent, Santa Clara, CA, USA) was used to measure the oxygen consumption rate (OCR) which relates to mitochondrial of live lymphocytes cells. Briefly, T cells were electroporated with CD19 CAR, IGK CAR, Kz-19BB CAR, 19z-KBB. Approximately 16 hours after electroporation, cells were seeded onto either Cell-Tak (Corning Inc, Corning, NY, US), anti-CD3 (OKT3) (eBiosciences, Thermo Fisher Scientific) or IgG coated 96-well XF-PS plates (Agilent, CA, USA). The density of cells determined to be 1×10^5^ cells/well in DMEM (ThermoFisher Scientific) XF unbuffered assay media, supplemented with 2 mM sodium pyruvate (Sigma-Aldrich, Norway), 10 mM glucose (Sigma-Aldrich), 2 mM L-glutamine (Thermo Fisher Scientific), adjusted to physiological pH (7.6). The cells were incubated in the absence of CO_2_ for one hour prior to Seahorse measurements (6 replicates per experiment). Initially, the cell basal respiration was measured for all groups. Next, oligomycin (Sigma-Aldrich), a potent F1F0 ATPase inhibitor (1 μM), was added and the resulting OCR was used to derive ATP production by respiration. Then, 1 μM of carbonyl cyanide p-trifluoromethoxyphenylhydrazon (FCCP) (Sigma-Aldrich) was injected to uncouple the mitochondrial electron transport from ATP synthesis, thus allowing the electron transport chain (ECT) to function at its maximal rate. The maximal respiration capacity was derived from each group by subtracting non-mitochondrial respiration from the FCCP measurement. Lastly, a mixture of antimycin A (Sigma-Aldrich) and rotenone (Sigma-Aldrich) was added, at 1 μM, to completely inhibit the electron transport and hence respiration, revealing the non-mitochondrial respiration.

### Multicellular tumor spheroid (MCTS) formation

Wells of a 96-well plate were coated with 50 μL of a 1.5% (w/w) solution of Agarose (Corning, New York, NY, USA) in PBS and left to polymerize during 60 min at room temperature. 1×10^3^ BL-41 or Granta-519 cells in 200 μL of complete RPMI-1640 was then added per well. BL-41 and Granta plates were spun down for 15 min at 1000g and incubated for 1.5 and 9.5 days at 37°C and 5% CO2, respectively. For the whole duration of the incubation, the plates were placed in an Incucyte S3 (Essen Bioscience Ltd., Newark, UK) with the following settings: 12 images/day, 1 image/well, 3 channels (phase, green and red). 50 μL of a 1:200 solution of Annexin V red (Essen Biosciences, UK) diluted in complete RPMI 1640 was added per well and the plate was consecutively incubated at 37°C, 5% CO2 for 15 min. Mock, IGK CAR, Kz-19BB and CD19 CAR transduced T cells previously washed and resuspended in complete RPMI-1640 medium were introduced in each well at a final concentration of 1×10^4^ cells/mL (50 μL/well). The plate was then put into an Incucyte S3 with the same settings as described above. Analysis of cytotoxicity was performed using Incucyte software.

### Statistical analysis

Student’s t-test, one-way or two-way ANOVA was used in the comparison of two groups. Half-maximal lysis was calculated by non-linear regression (curve fit) on *GraphPad Prism*^®^ (GraphPad Software, Inc.).

## Data Availability

The data that support the findings of this study are available from the corresponding author upon reasonable request.

## Acknowledgments

We are grateful to Anne Fåne, Marit Renée Myhre and Elizabeth Baken for expert technical assistance. We thank the Flow Cytometry Core Facility and The Core Facility for Advanced Light Microscopy at the Institute for Cancer Research, Oslo University Hospital.

## Author Contributions

SW, HK, EMI, JHM designed the experiments, HK, PD, EB, AJ performed the experiments, SW, HK, EMI, PD analyzed the experiments, EBS, GK provided essential material and support, HK, SW, EMI and PD wrote the text.

## Abbreviations and nomenclature

BCR: B cell receptor
BLI: Bioluminescence
CD3ζ: CD3 zeta
CAR: chimeric antigen receptor
Ig: immunoglobulin
IGK: immunoglobulin kappa
IGL: immunoglobulin lambda
*in vitro*: transcribed messenger
IVT mRNA: RNA
κ: kappa
λ: lambda
scFv: single chain variable fragment
sIgG: soluble IgG
TCR: T cell receptor

---

**Supporting Figure 1:**
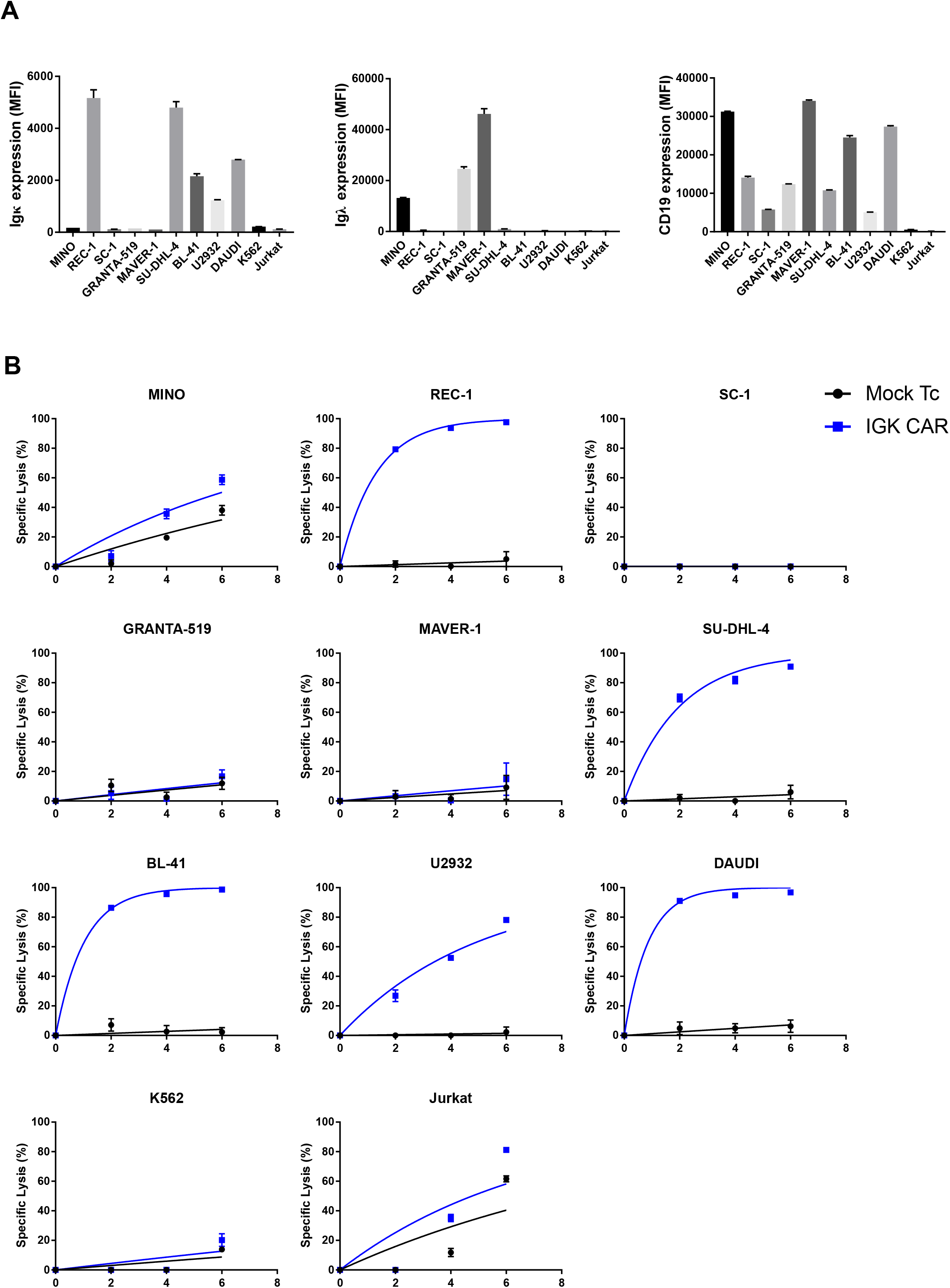
IGK CAR-expressing T cells target and specifically eliminate Igκ+ target cells. (A) Expression levels of Igκ, Igλ light chains and CD19 receptor for each cell lines. Data represent mean ± S.D. of duplicates. (B) BLI assay performed by Mock T cells and IGK CAR electroporated T cells at an E:T ratio of 25:1 for 6 hours. Data represent mean ± S.D. of quadruplicates. Representative data from one of two independent experiments are shown.

**Supporting Figure 2:**
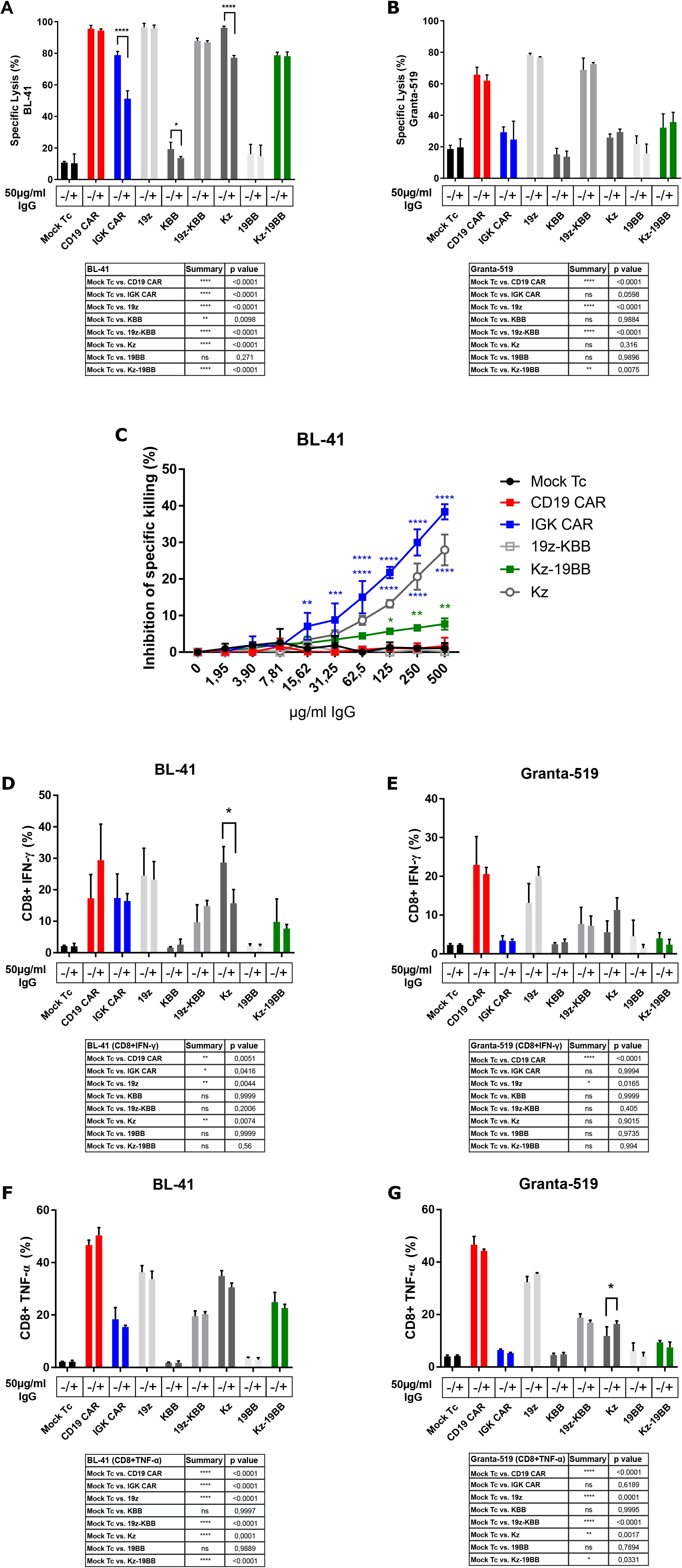
Combinatorial CAR T cells are less sensitive to serum inhibition compared to individual components. (A-B) BLI killing assay of Mock T cells and CAR construct electroporated T cells co-cultured with Igκ^+^ BL-41 or Igκ^-^ Granta-519 cell lines for 10 hours (10:1 E:T ratio) in the presence or absence of serum purified IgG (50μg/ml). Data represent mean ± S.D. of quadruplicates. Representative data from one of three independent experiments are shown. (C) BLI killing assay of Mock T cells and CAR construct electroporated T cells co-cultured with BL-41 cell line for 10 hours in presence of increasing concentration of IgG (10:1 E:T ratio). Specific lysis inhibition corresponds to the difference of cytotoxic capacity of each construct between IgG+ and IgG-conditions. Data represent mean ± S.D. of triplicates. One-way ANOVA was performed between Mock T cell and other groups for each IgG concentration. (D-G) CD8+ T-cell intracellular cytokine staining (IFN-γ and TNF-α) after co-cultivation with BL-41 or Granta-519 cell lines in the presence or absence of IgG at 50μg/ml (for 24 hours, 1:2 E:T ratio) Data represent mean ± S.D. of triplicates. Representative data from one of two independent experiments are shown. (A,B,D-G) Significance was assessed by Student t-test comparing IgG+ and IgG-conditions and by one-way ANOVA between Mock T cells and any other group (IgG-). \**P* < 0.05, \*\**P* < 0.01, \*\*\**P* < 0.001, ****P<0.0001.

**Supporting Figure 3:**
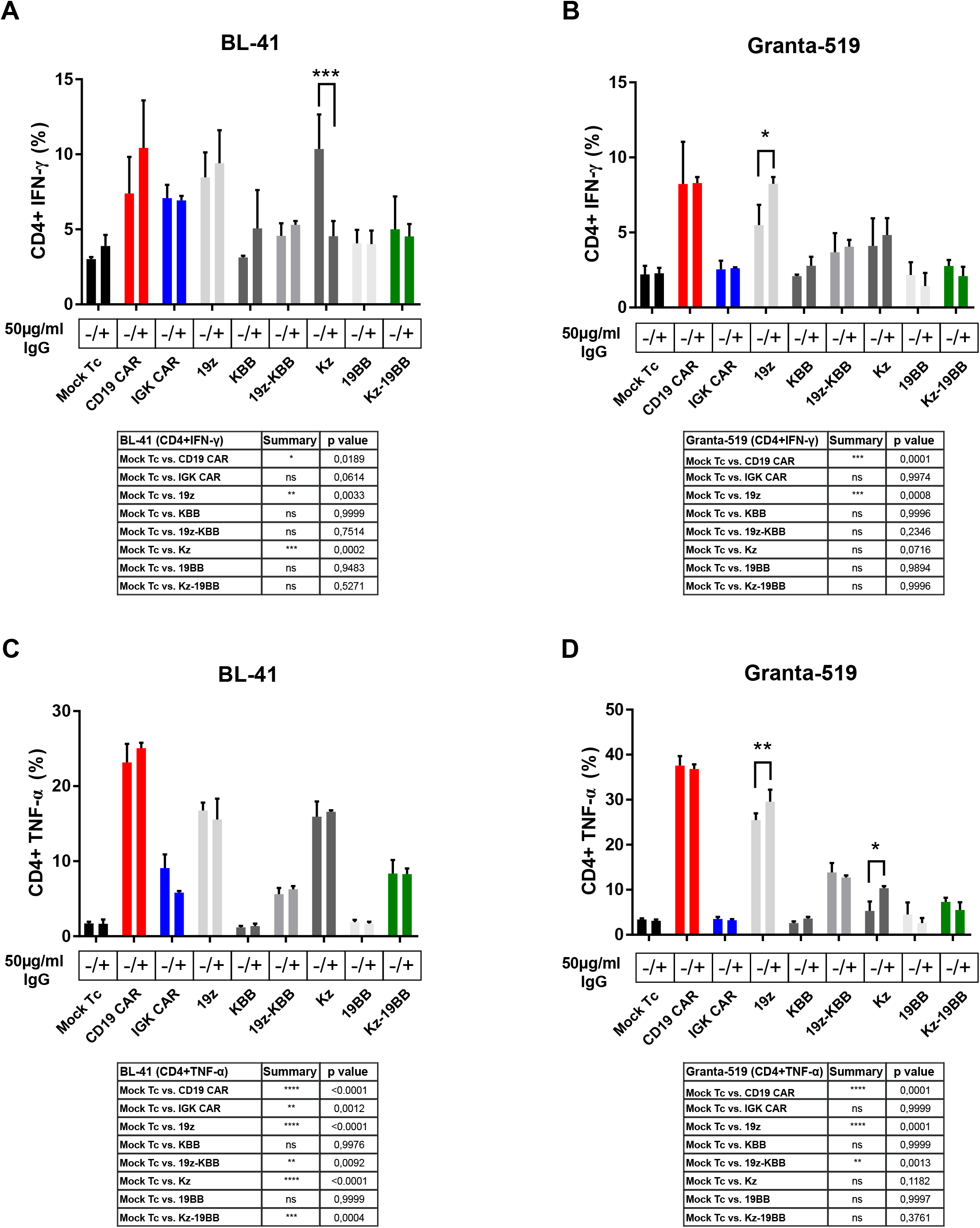
Combinatorial CARs limit serum inhibition while preserving CD3ζ driven specificity. (A-D) CD4+ T-cells intracellular cytokine staining (IFN-γ and TNF-α) after co-cultivation with BL-41 or Granta-519 cell lines in the presence or absence of IgG (50μg/ml) (for 24 hours, 1:2 E:T ratio) Data represent mean ± S.D. of triplicates. Representative data from one of two independent experiments are shown.

**Supporting Figure 4:**
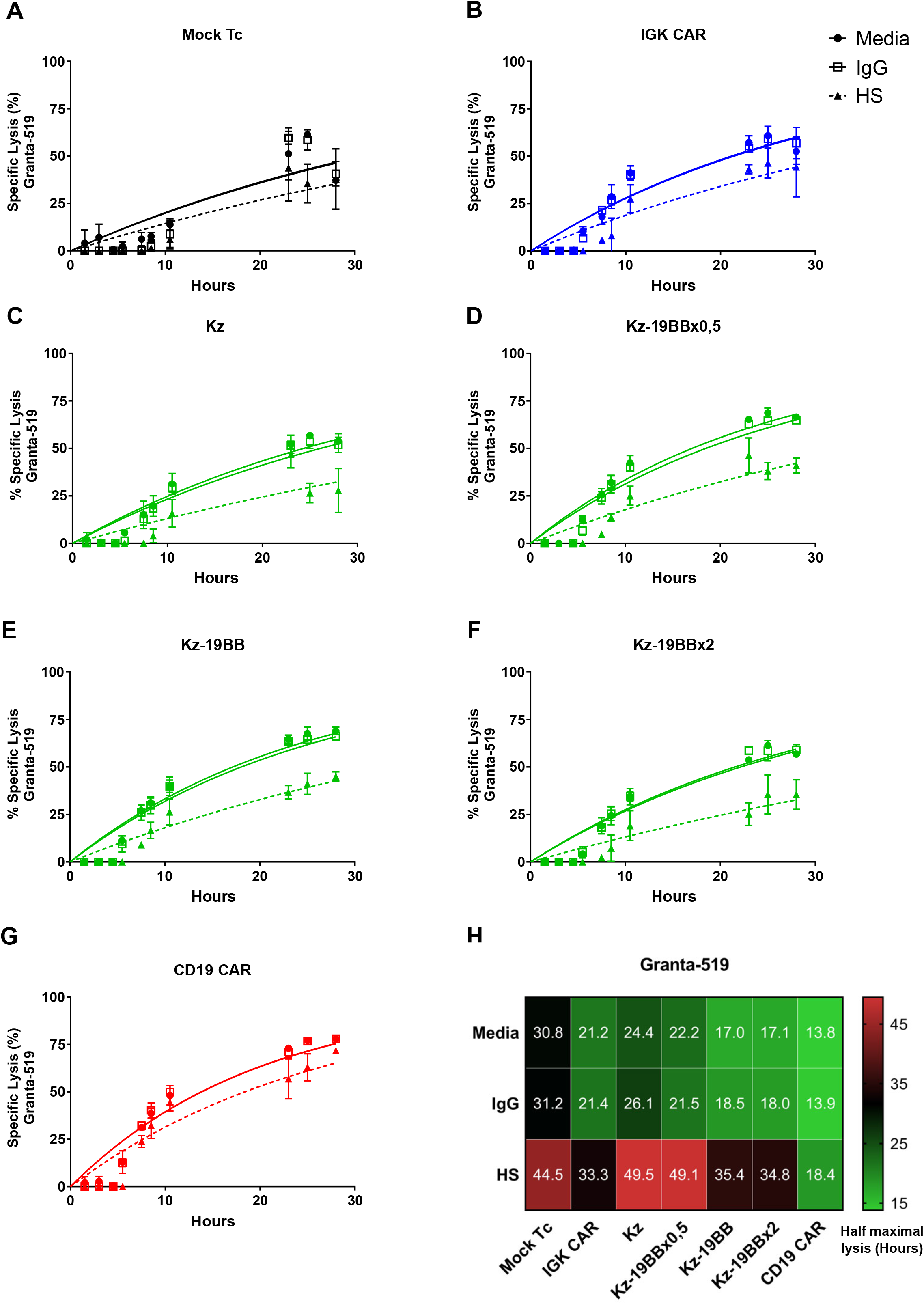
Kz-19BB expressing T cells also target CD19+ cells with each increased concentration of 19BB part. (A-G) BLI killing assay of Mock and CARs construct mRNA electroporated T cells co-cultured with Granta-519 cell lines either in presence or absence of IgG (50μg/ml) or 25% human serum. H) Heat maps of half maximal lysis values obtained via one phase exponential fitting against Granta-519. Data represent mean ± S.D. of quadruplicates. Representative data from one of three independent experiments are shown. \**P* < 0.05, \*\**P* < 0.01, \*\*\**P* < 0.001, ****P<0.0001.

**Supporting Figure 5:**
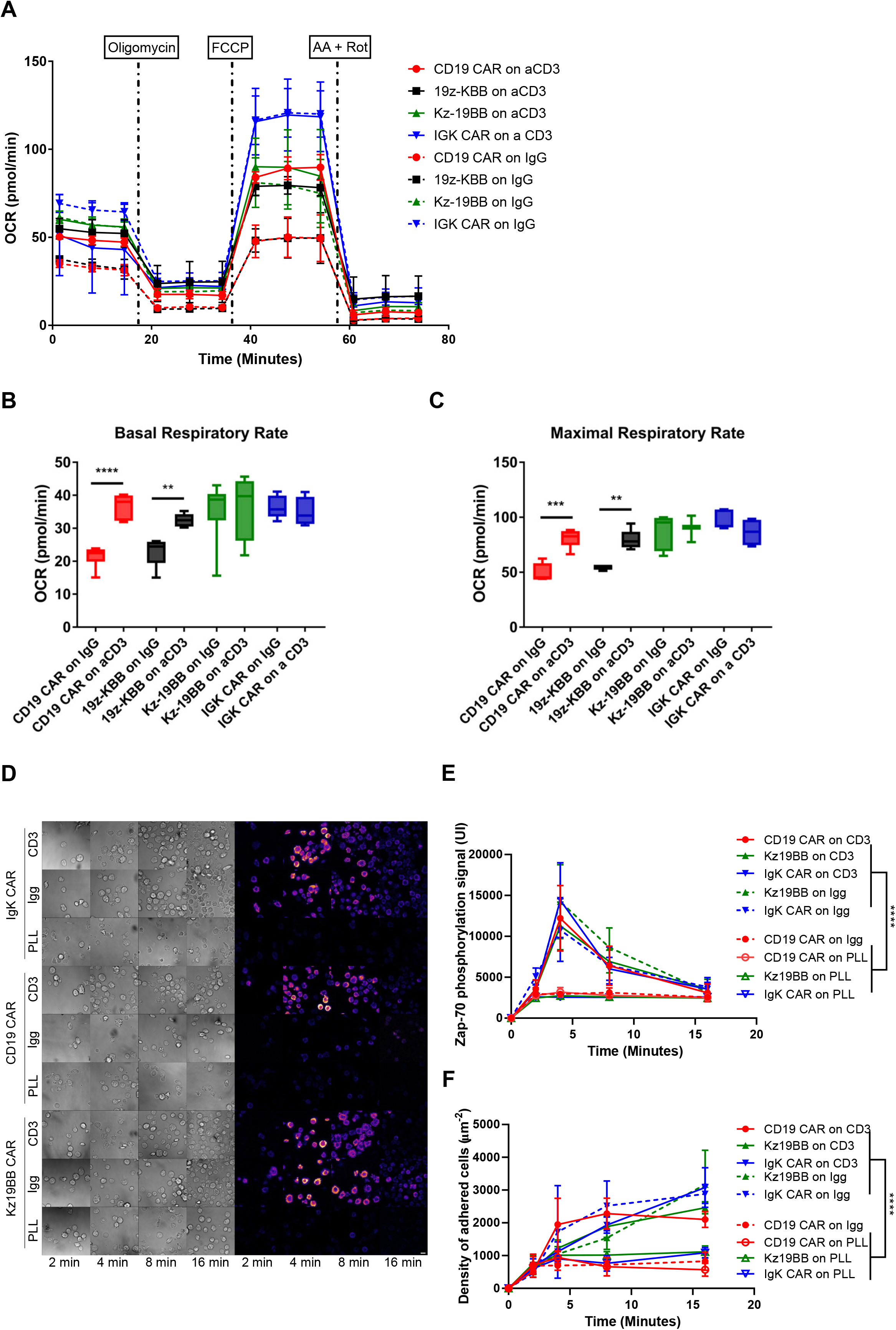
Combinatorial CAR, Kz-19BB maintains IGK CAR properties and is activated in presence of IgG. (A) Primary T cells transfected with CD19 CAR, 19z-KBB, Kz-19BB or IGK CAR coding mRNA were incubated on either anti-CD3 or serum purified IgG coated wells 18 hours after electroporation. Oxygen consumption rate (OCR) was assessed via a Seahorse assay. (B-C) Basal and maximal respiratory rates for each condition. Data represent mean ± S.D. of hexaplicates. Significance assessed by Student’s t-test. (D) Kz-19BB, CD19 CAR or IGK CAR constructs electroporated T cells were incubated on anti-CD3, IgG or Poly-L-lysine (PLL) for 2, 4, 8 or 16 minutes. Cells were stained with anti-phospho-ZAP70-PE. Representative results for each time point acquired by confocal and corresponding bright field images are shown. Scale bar represents 5μm. (E) ZAP70 phosphorylation signal intensity as analyzed from micrographs. Significance between each construct’s anti-CD3 and IgG response relative to their PLL response was assessed by two-way ANOVA with Tukey’s correction (n = 50). (F) Surface adherence properties of each construct when in contact with anti-CD3 and IgG were assessed relative to their PLL response by mixed-effects model (REML). \*\**P* < 0.01, \*\*\**P* < 0.001, ****P<0.0001.

**Supporting Figure 6:**
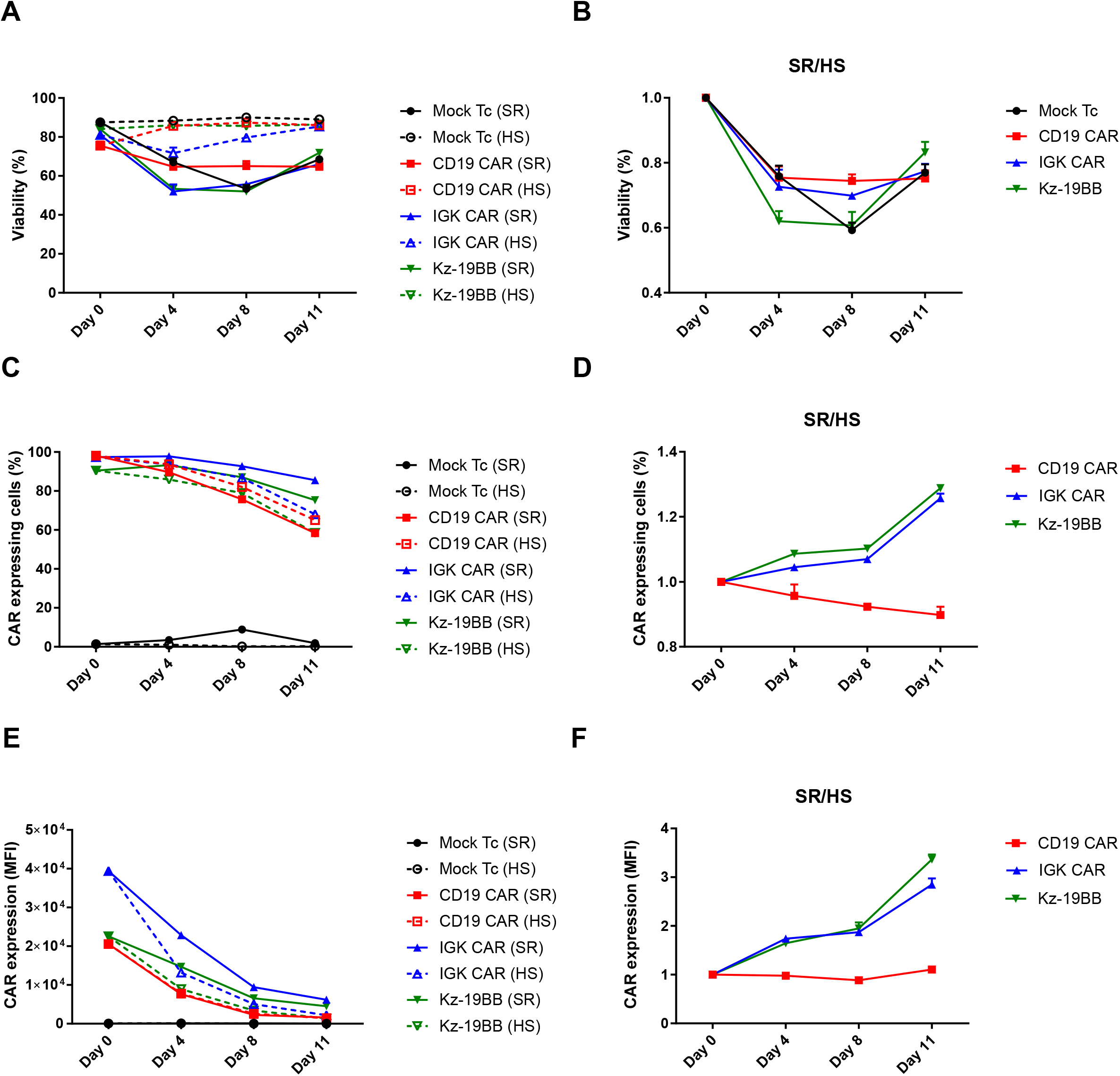
Stable IGK and Kz-19BB CAR-expressing T cells downregulate CAR expression when expanded in human serum. (A-F) Mock and CAR expressing T cells were expanded in either SR (serum replacement) or HS (human serum) containing X-VIVO-15 complete media for 11 days. Cell viability and other properties were evaluated on days 0, 4, 8 and 11. (A-B) Cell viability is monitored with a cell counter utilizing trypan blue exclusion. SR values are divided over HS values to observe the general trend between each expansion method. (C-F) CAR expressions were assesed as before and plotted either as percentage or mean fluorescent intensity (MFI). Similarly, SR values are divided over HS to observe a general trend. Data represent mean ± S.D. of triplicates. Representative data from one of three experiments are shown.

**Supporting Figure 7:**
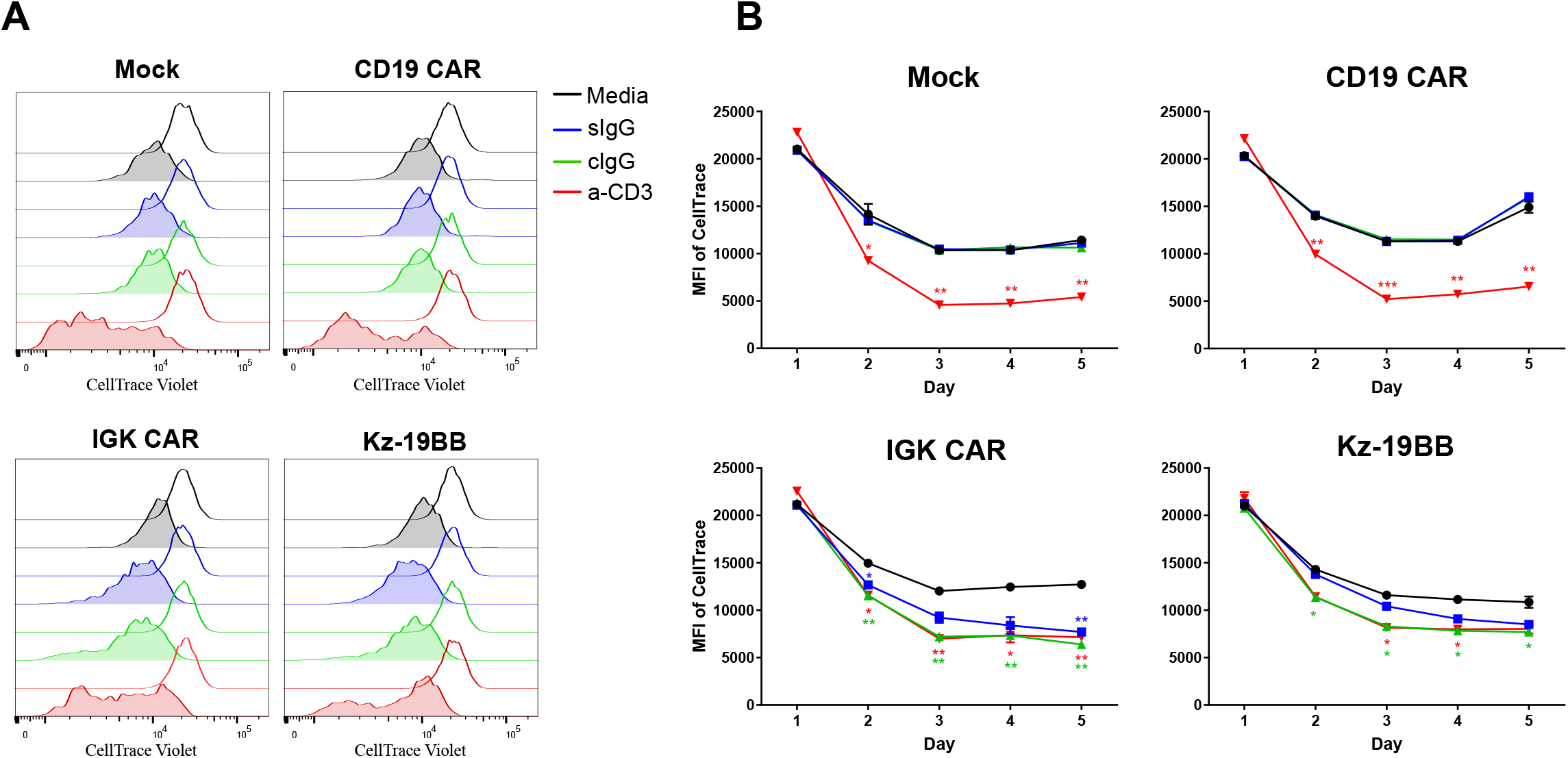
IGK CAR and Kz-19BB mount a proliferative immune response to coated IgG as well as soluble. (A) Mock or construct transduced T cells were cultured in the presence of media or sIgG or on anti-CD19 or IgG coated wells for 5 days. Prior to the experiment T cells were stained with CellTrace Violet and proliferation status was assessed every day through flow cytometry. Representative results show the difference between each group’s Day 1 (line) and Day 5 (filled) status. (B) Proliferation assessment of each T cells group over the course of 5 days. Statistical test was performed between each group’s media and other conditions by student’s t-test. Data represent mean ± S.D. of duplicates. \**P* < 0.05, \*\**P* < 0.01, \*\*\**P* < 0.001, ****P<0.0001.

**Supporting Figure 8:**
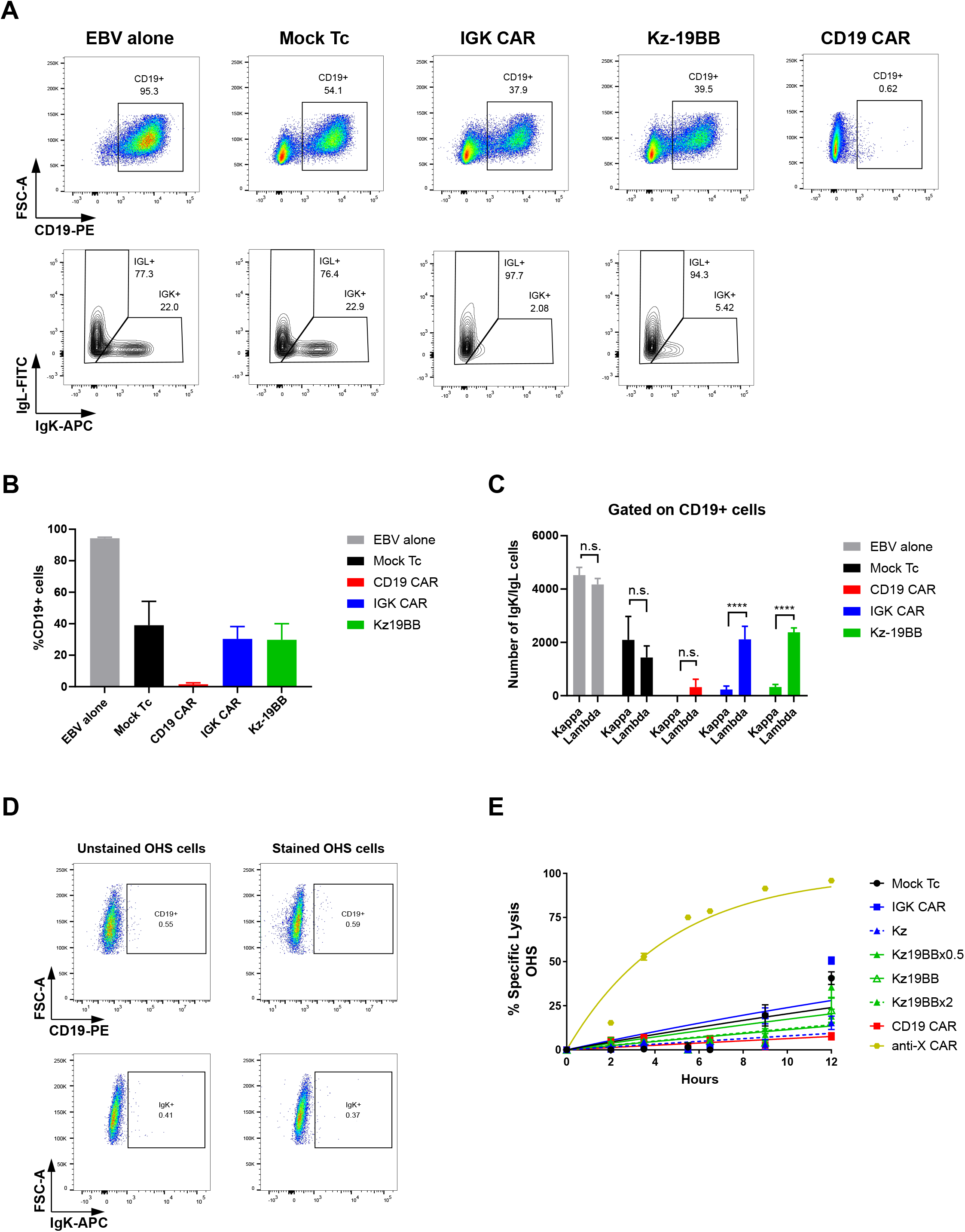
Combinatorial CAR, Kz-19BB selectively eliminates Igκ+ EBV cells in a mixed environment. (A-C) Retrovirally transduced T cells co-cultured for 12 hours with both BL-41 and Granta-519 target cell lines at a ratio of 2:1:1, respectively. After co-culture cells were stained with anti-CD19-PE, anti-Igκ APC and anti-Igλ-FITC. Data represent mean ± S.D. of quadruplicates. Representative data from one of three experiments are shown. Significance was assessed by Student t-test between kappa+ and lambda+ cell numbers. \**P* < 0.05, \*\**P* < 0.01, \*\*\**P* < 0.001, ****P<0.0001. (D) OHS cell line was stained with anti-CD19-PE and anti-Igκ APC. (E) BLI killing assay of Mock and CARs construct electroporated primary T cells co-cultured with OHS cell line at an E:T ratio of 10:1 for 12 hours. Data represent mean ± S.D. of triplicates. Representative data from one of two experiments are shown.

